# Replicating without stress: a dengue replicon model reveals distinct host rewiring to accommodate persistence

**DOI:** 10.1101/2025.09.27.678963

**Authors:** Karen W. Cheng, Damian Kim, Frank McCarthy, Kuo-Feng Weng, Leah Dorman, Joshua E. Elias, Amy Kistler, Ranen Aviner

## Abstract

Self-replicating viral RNAs, or replicons, are widely used to study virus-host interactions and screen antiviral compounds, yet their ability to model persistent infection and host adaptation remains poorly understood. Here, we characterize a stable subgenomic dengue virus serotype 2 (DENV2) replicon (DENV2-rep) that supports long-term viral RNA replication in human hepatoma cells without triggering cytotoxicity or canonical stress responses. Compared to live DENV2 infection, which induces ER stress and global translation shutdown, DENV2-rep cells maintain active translation and polysome integrity, while mounting an antiviral transcriptional response and resisting superinfection. Transcriptomic and proteomic profiling reveals that DENV2-rep cells accommodate long-term replication by remodeling of metabolic, secretory, and cytoskeletal pathways. In contrast, acute infection uncouples transcription from protein synthesis, limiting host proteome remodeling. We also detected marked depletion of the NS5 RNA polymerase in the DENV2-rep context. Together, these findings highlight host pathways that can be rewired to support persistent RNA replication and uncover post-translational regulation of viral proteins as a potential vulnerability. Our work clarifies the strengths and limitations of replicons as models for infection and reveals how persistent vRNA replication is tolerated through selective remodeling of translation, organelle composition, and protein stability.

## Introduction

Dengue virus (DENV), a mosquito-borne flavivirus, is a major global health threat responsible for over 100 million new infections annually[1]. Despite decades of research, effective antiviral therapies remain elusive. Other than targeting Dengue structural and nonstructural proteins[2,3], another promising strategy involves targeting host enzymes that are essential for viral replication but dispensable for host viability (so-called host-directed therapies [4]). However, realizing this strategy requires a deeper understanding of how DENV interacts with and reprograms the host cellular environment.

DENV has a positive strand RNA genome of about 11 kilobase (kb) in length. The genome is 5’ capped, lacks a poly-A tail, and translated into the ER membrane as a single long polyprotein, which is cleaved by host and viral proteases to release viral proteins into the cytosol and ER lumen[5]. This process places high biosynthetic demands on the ER and remodels its architecture[6]; therefore, acute DENV infection commonly triggers a canonical stress response that involves activation of the unfolded protein response (UPR)[7,8], global translation inhibition[9,10], and suppression of innate immune signaling [11,12]. Although the UPR typically functions to restore homeostasis or eliminate stressed cells, many viruses, including DENV, co-opt it to promote replication and evade immune detection[13].

In contrast to acute cytopathic infections in human cells, DENV establishes long-term, noncytopathic infection in mosquito vectors, where viral RNA (vRNA) and protein synthesis persist without triggering cell death[14]. Persistent DENV infection was also observed in immunocompromised patients[15]. While differences in host species, ER stress tolerance, and immune response likely contribute to this dichotomy, the molecular mechanisms that enable persistence in some contexts and cytotoxicity in others remain poorly understood. Replicons—engineered self-replicating vRNAs that express nonstructural proteins but lack structural genes—offer a powerful platform to investigate this question[16,17]. By isolating the effects of vRNA translation and replication from other lifecycle steps e.g. entry, assembly, and egress, replicons enable the study of how cells respond to viral replication without the confounding effects of virion production.

Although DENV replicons have been used to screen for antivirals and study RNA replication and replication complex dynamics, most commonly to model acute short-term infection[16–19], it remains unclear whether host responses to replicons differ from live infection. Moreover, most replicon studies rely on transient transfection, capturing acute responses rather than long-term adaptation[16,17]. This raises fundamental questions about how the host transcriptome, proteome, and stress signaling pathways are remodeled to accommodate persistent vRNA replication in the absence of structural proteins.

Here, we use a stable DENV2 replicon to systematically compare host responses to acute and persistent DENV2 replication in human hepatoma cells. Using transcriptomics, proteomics, and biochemical assays, we uncover differences in translation, stress responses, and protein turnover. Replicon cells avoid canonical ER stress and translation shutdown seen in acute infection, instead maintaining active protein synthesis and mounting a sustained antiviral response. Integrated analysis reveals that mRNA and protein changes are tightly coordinated in replicon cells, particularly across membrane-bound organelles and metabolic pathways, while acute infection leads to uncoupling of mRNA and protein levels. We further identify rapid turnover of the viral polymerase NS5 in replicon cells, highlighting post-translational regulation of protein stability as a potential control point for persistent replication. Together, these findings define the host cell state that accommodates persistent vRNA replication and demonstrate the utility of replicons for dissecting long-term virus-host interactions.

## Results

### Cells stably expressing DENV replicon support persistent RNA replication without cytotoxicity

DENV2 genome encodes both structural and nonstructural proteins, enabling the full viral life cycle from entry and uncoating, through RNA replication and translation, to virion assembly and egress (Fig. 1a). In contrast, a recently developed stable DENV2 replicon model (DENV2-rep) [20] encodes only nonstructural (NS) proteins and therefore permits vRNA translation and replication without formation of infectious particles (Fig. 1b, left). In this subgenomic replicon, the structural genes (C-prM-E) were replaced with a sequence encoding destabilized EGFP and a selectable marker, separated from viral nonstructural proteins by an F2A ribosome skip site. To generate DENV-2 rep, in vitro transcribed replicon RNA was transfected into Huh7.5.1 cells followed by antibiotic selection (Fig. 1b, right and ED Fig. 1a). We hypothesized that the resulting stable line (DENV2-rep), originally developed for CRISPR screening[21], can serve as a platform to study long-term adaptations to persistent dengue RNA replication.

**Figure 1.**
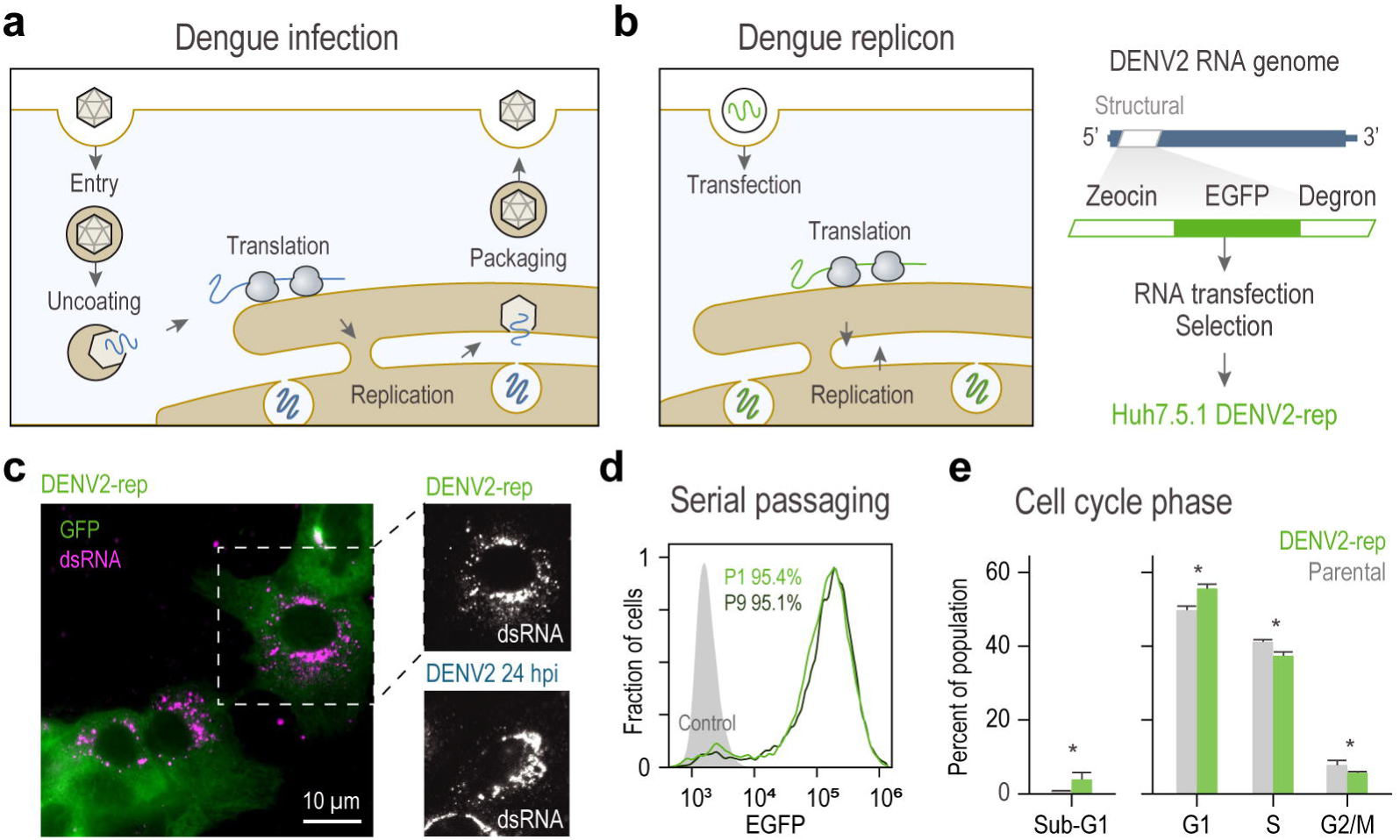
A stable DENV2 replicon supports persistent RNA replication without cytotoxicity. (a-b) Schematics of DENV2 life cycle (a) compared with the subgenomic DENV2 replicon (b). The replicon encodes nonstructural proteins NS1-NS5 flanked by native 5’ and 3’ untranslated regions. Structural genes were replaced with a cassette encoding a selectable Zeocin-eGFP fusion protein with a C-terminal PEST sequence (Degron), separated from downstream viral sequences by an F2A ribosome skip site. To generate a stable replicon line (DENV2-rep), Huh7.5.1 cells were transfected with in vitro transcribed replicon RNA, followed by antibiotic selection and expansion. (c) Immunofluorescence staining of double-stranded RNA (dsRNA) in fixed DENV2-rep cells and controls infected with DENV2 for 24 hours. hpi, hours post-infection. dsRNA accumulates in perinuclear puncta consistent with active replication. (d) Flow cytometry analysis of EGFP expression across 9 serial passages (P1-9). More than 94% of cells retained replicon expression after 9 passages, demonstrating stability over time. Shown are representative plots from n=3 repeats for each passage. (e) Cell cycle analysis of parental and replicon cells by DNA staining and flow cytometry. DENV2-rep cultures show modest increase in G1/Sub-G1 and reduction in S/G2-M compared to parental cells, indicating low cytotoxicity. Bar graphs show means ± s.d. of 3 independent repeats. *, two-tailed Student’s t test p-value = 0.017, 0.026, 0.050, 0.024 for sub-G1, G1, S and G2/M.

Immunofluorescence staining of fixed cells confirmed that replicon RNA produces the characteristic double-stranded RNA puncta seen in parental cells infected with DENV2 at a high multiplicity of infection (MOI), consistent with formation of replication compartments (Fig. 1c). Replication compartments were previously shown to form in the absence of structural proteins[18]. vRNA accumulated to similar levels under both conditions (ED Fig. 1b). Treatment with the flavivirus polymerase inhibitor MK-0608[22] reduced RNA levels (ED Fig. 1c), confirming that maintenance depends on ongoing RNA synthesis by viral replication complexes. Flow cytometry analysis demonstrated robust stability of replicon expression under antibiotic selection; more than 94% of cells retained EGFP expression across 9 serial passages (Fig. 1d).

Despite continuous RNA replication, DENV2-rep cultures remained healthy and showed no overt cytopathic effects. Cell-cycle analysis revealed only a modest shift—slightly more cells in G1/Sub-G1 and fewer in S and G2/M—compared to parental Huh7.5.1 cells (Fig. 1e and ED Fig. 1d). Together, these data establish that dengue replicon RNA can be stably maintained in human hepatoma cells, supporting persistent replication without triggering cytotoxicity.

### DENV2-rep maintains high viral RNA load with adaptive sequence change

We next performed RNA-seq on DENV2-rep cells and on parental Huh7.5.1 cells infected with DENV2 at a high MOI, with uninfected controls (Fig. 2a). Infected cells were collected at 24 hours post-infection (hpi) to match vRNA levels between conditions (ED Fig. 1a), capture host transcriptional responses e.g. ER stress that are first detected in Huh7-derived cells at 24 hpi[11], and avoid the cytopathic effects that dominate at 48 hpi[23]. As expected, vRNA was detected at high levels in both conditions; about 8-and 20-fold higher than GAPDH mRNA, amounting to ∼2 and 6% of total coding transcripts in DENV2-rep and infected cells, respectively (Fig. 2a and Table S1). Genome coverage was mostly similar in both conditions, confirming sustained replication of the full-length vRNA without major deletions (Fig. 2b). DENV2-rep cells had higher levels of RNA fragments mapping to the 3’ untranslated region (UTR) (Fig. 2c). DENV2 3’ UTR fragments, otherwise known as subgenomic flaviviral RNA (sfRNA), are generated during infection by cellular exonucleases and increase vRNA replication by suppressing interferon production[24]. DENV sfRNAs accumulate to higher levels in persistently infected mosquito cells[25].

**Figure 2.**
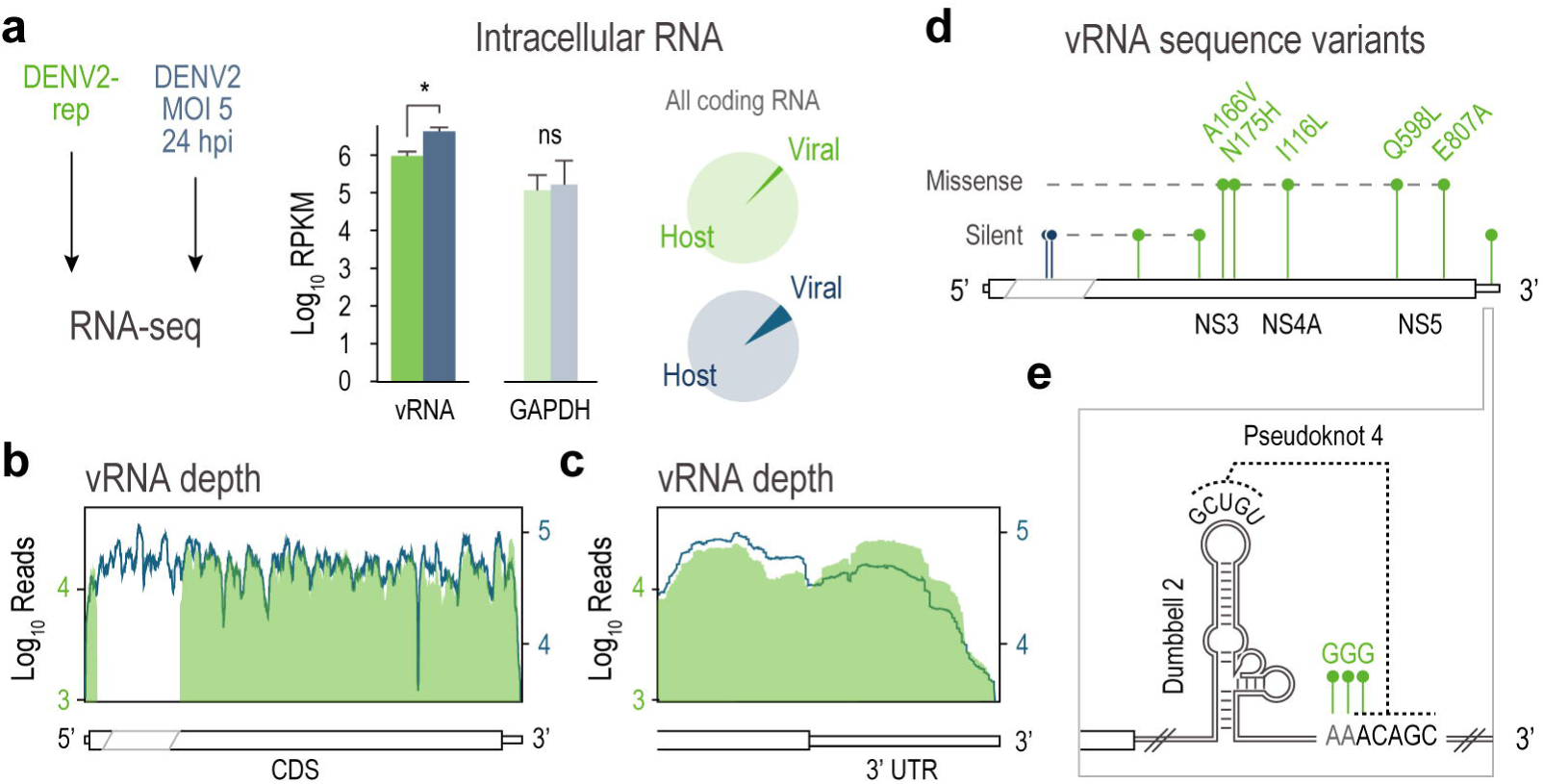
DENV2-rep maintains high viral RNA levels with adaptive sequence changes. (a) RNA-seq of total intracellular RNA detects comparable levels of vRNA in DENV2-rep and parental cells infected for 24 hpi. MOI, multiplicity of infection. Bar graphs show means ± s.d. of 3 independent repeats. *, two-tailed Student’s p-value < 0.01. Pie charts show that vRNA reads account for 1.7±0.13 and 5.9±0.61% of all coding RNA reads in replicon and infected cells, respectively. Means ± s.d. (b-c) Sequencing depth plots showing reads mapping to full-length vRNA (b) and reads mapping to the 3’ end of the viral genome (c). (d) Variant analysis detected two synonymous mutations in infection (blue) and multiple synonymous and nonsynonymous mutations in replicon cells (green). Allele frequency for all variants shown is >95% in all 3 repeats. (e) Substitutions in the replicon 3′ UTR map to the conserved pseudoknot 4, formed between dumbbell 2 (DB2) and the indicated unstructured sequence.

Analysis of single nucleotide variants in vRNA (Table S2) revealed only two synonymous mutations in acutely infected cells, which mapped to the structural coding region (Fig. 2b, blue, and ED Fig. 2a). One of these mutations was previously reported to arise in Huh7 cells [26]. In contrast, vRNA in DENV2-rep accumulated two silent and five missense mutations in the coding sequence, and three substitutions in the 3′ UTR (Fig. 2d and ED 2a). Two missense mutations in the RNA-dependent RNA polymerase NS5 mapped to motif B and the priming loop, which facilitate RNA sliding and primer-independent replication[27]. Based on RNA structure simulations, these 3′ UTR substitutions likely destabilize a conserved structural element (dumbbell 2/pseudoknot 4 [DB2/PK4]) (Fig. 2e and ED Fig. 2b-c). Structure destabilizing mutations in this domain occur with passaging in Huh7 cells[28], improve long-term replication in mosquitos[29], and reduce cytotoxicity in hamster and monkey cells[29]. All mutations were present at high allele frequencies (>95%), in all three replicates, suggesting adaptive selection during long-term replication (Table S2).

### Persistent replication activates antiviral programs without ER stress

At the host level, transcriptional responses diverged between replicon and acute infection. DENV2-rep cells showed far broader changes, with 1,684 differentially expressed genes (DEGs; fold change > 2, FDR < 0.05) versus only 47 in live infection (Fig. 3a). This difference likely reflects both the long-term adaptation of replicon cells to vRNA replication and the asynchronous, heterogeneous nature of acute infection, even at a high MOI, which dilutes transcriptional signals in bulk analyses[11]. The homogeneity of antibiotic-selected replicon cells may thus reveal host programs that are harder to detect during cytopathic infection.

**Figure 3.**
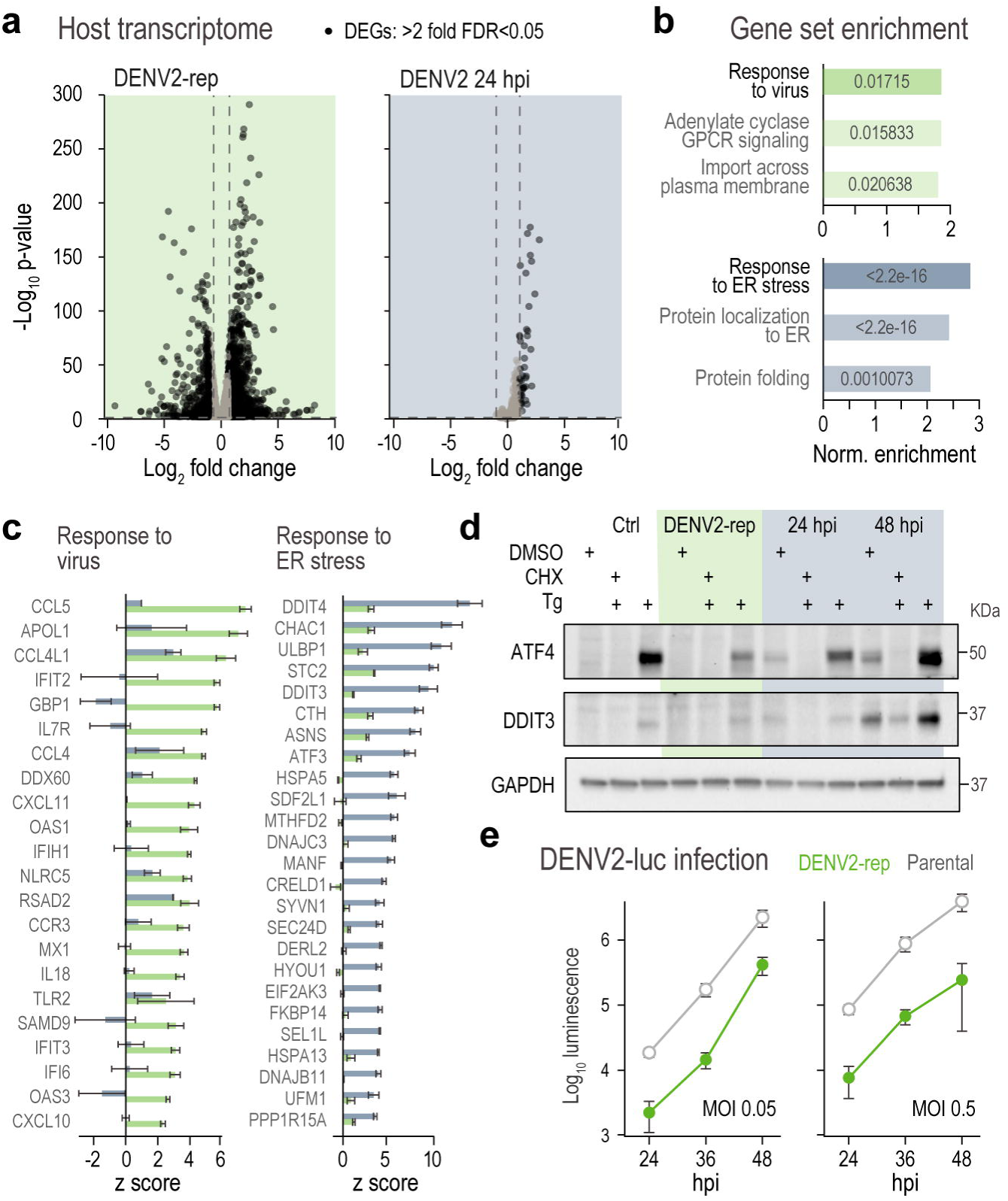
Persistent replication in DENV2-rep activates antiviral programs without ER stress. (a) Differentially expressed genes (DEGs) in RNA-seq data from replicon and infection relative to uninfected parental controls. Replicon cells show 1,684 DEGs versus 47 in infection (FDR < 0.05, FC >2). (b) Gene set enrichment analysis (GSEA) comparing replicon and infection. Virus-response pathways are enriched in replicon cells, whereas ER stress pathways dominate in infection. (c) Bar graphs showing top antiviral genes induced in DENV2-rep cells and top ER stress genes induced in infection. Means ± s.d. of 3 independent repeats. (d) Immunoblots for ATF4 and DDIT3 (CHOP). DENV2 infection induces UPR effectors in a time-dependent manner, whereas replicon cells lack induction but respond to chemical ER stress. Cells were treated with either 10 µg cycloheximide (CHX) or 1 µM thapsigargin (Tg) for 4 hours before harvesting. Blots are representative of 3 independent repeats. (e) DENV2-rep and parental controls were infected with infectious DENV2 that expresses renilla luciferase from the viral genome. Luminescence was measured at the indicated times. Shown are means ± s.d. of 3 independent repeats.

To account for bulk dilution effects, we applied gene set enrichment analysis (GSEA), which captures pathway-level trends rather than absolute fold changes. GSEA revealed strikingly distinct stress responses: DENV2-rep cells enriched for virus response, whereas infected cells showed the hallmark ER stress response (Fig. 3b and ED Fig. 3a-b). Antiviral responses in DENV2-rep included interferon-stimulated genes (ISGs) e.g. OASL, IFIT2, MX1, RSAD2, and CXCL10, which were not induced in acute infection (Fig. 3c and ED 3c), consistent with innate immune suppression[11]. However, similar ISG signatures were detected in dengue patients and vaccine recipients[30,31] suggesting that the replicon may capture some aspects of *in vivo* host responses. ER stress response genes induced by acute infection included HSPA5 (BiP), DDIT3 (CHOP), EIF2AK3 (PERK), and PPP1R15A (GADD34) [32], which were not induced in DENV2-rep cells (Fig. 3c and ED 3c). This supports the hypothesis that Dengue structural proteins are the main drivers of PERK/ATF4-mediated UPR during infection[33].

To validate these transcript-level findings, we measured protein levels of ATF4 and DDIT3, downstream effectors of ER stress[32], by immunoblotting. Live DENV2 infection induced both proteins in a time-dependent manner, consistent with stress-driven activation of the UPR (Fig. 3d). Cycloheximide treatment to stop further production of viral proteins reduced ATF4 and DDIT3 to background levels. In contrast, DENV2-rep cells showed no detectable ATF4 or DDIT3 at baseline but upregulated them in response to chemical ER stress, confirming they are UPR competent.

Finally, to test whether the elevated antiviral response in DENV2-rep cells confers functional protection, we challenged them with infection with a luciferase-expressing DENV2 strain (DENV2-Luc)[26]. DENV2-rep cells produced significantly lower levels of luciferase (≥1 log reduction at all timepoints), consistent with reduced replication of DENV2-Luc (Fig. 3e). Together, these findings reveal that persistent DENV2 RNA replication triggers a distinct transcriptional response that activates antiviral defenses while avoiding canonical ER stress, establishing a steady-state host environment that is permissive but noncytopathic.

### DENV2-rep cells maintain translation and remodel the host proteome

DENV2 is known to suppress host translation through both ER stress and direct modulation of the translation apparatus [10,34,35]. We next asked whether persistent replication in DENV2-rep cells alters translation levels and, as a result, proteome composition. We first compared the distribution of active ribosomes across conditions using sucrose gradient fractionation and polysome profiling. As expected, live DENV2 infection markedly reduced polysome abundance and caused a shift toward 80S monosomes, consistent with a global translation block (Fig. 4a). In contrast, DENV2-rep cells maintained intact polysomes resembling uninfected controls, indicating that translation remains high despite ongoing vRNA replication.

**Figure 4.**
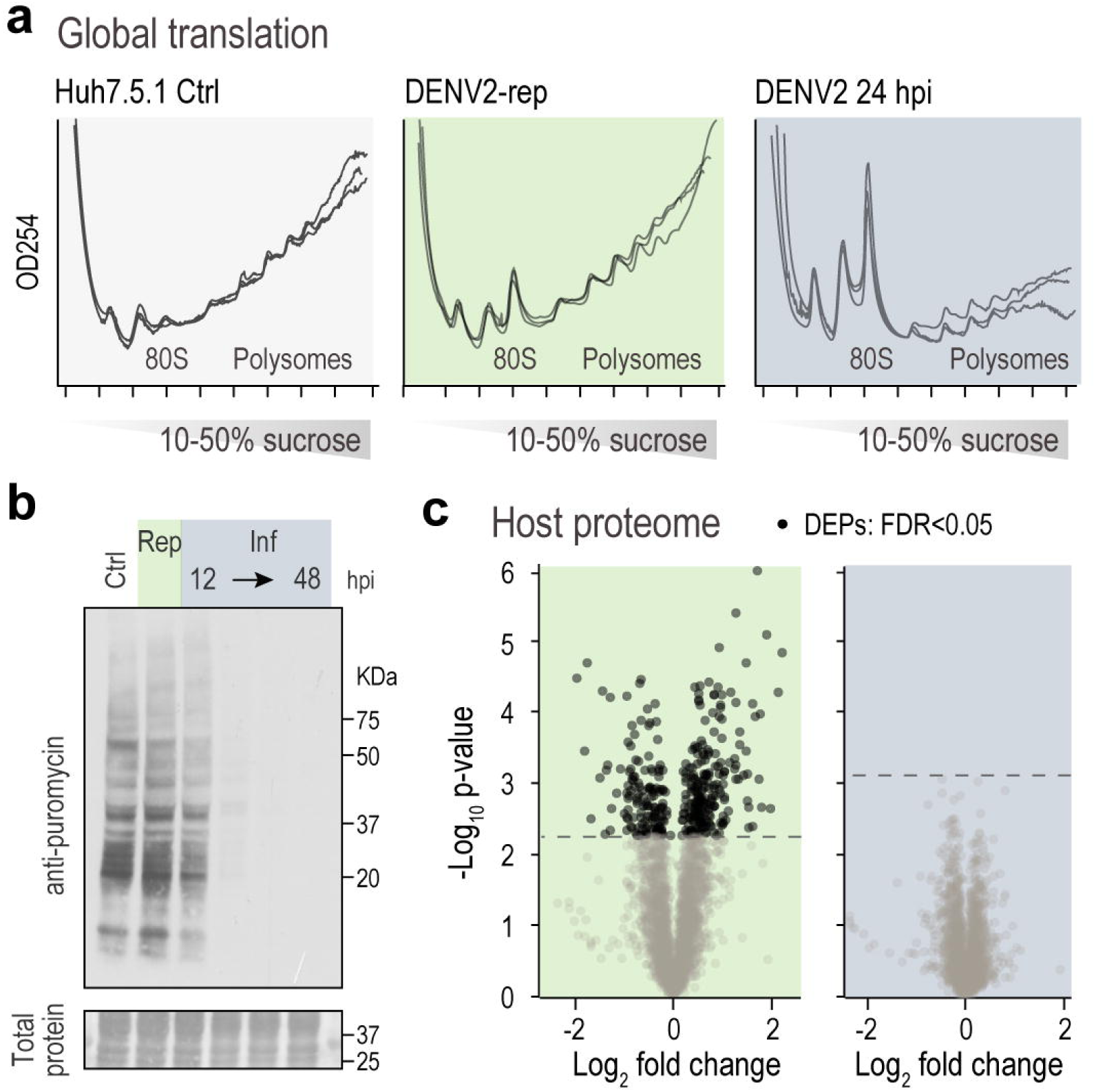
DENV2-rep cells maintain active translation in contrast to acute infection. (a) Polysome profiles from sucrose gradient fractionation. DENV2 infection shifts ribosomes toward 80S monosomes, indicating global translation repression, whereas replicon cells retain intact polysomes similar to controls. Shown are overlaid traces from 3 independent repeats. (b) Puromycin incorporation assay. Cells were treated with 1 µM puromycin for 15 min before harvesting. Nascent peptide synthesis, detected by an anti-puromycin immunoblot, is reduced in infected cells beginning at 12-24 hpi but preserved in replicon cells. Blots are representative of 2 independent repeats. (c) Differentially expressed proteins (DEPs) in mass spectrometry data from replicon and infection relative to uninfected parental controls. Replicon cells show 228 DEPs versus none in infection (FDR < 0.05).

Next, we used puromycin labeling to measure nascent polypeptide production. Consistent with translational repression, puromycin incorporation was significantly reduced in DENV2-infected cells beginning between 12 and 24 hpi (Fig. 4b). In contrast, DENV2-rep cells maintained robust puromycin labeling comparable to controls, confirming preserved translation capacity.

To determine how differences in translation affect proteome dynamics, we analyzed matched samples from DENV2-rep and DENV2-infected cells by label-free quantitative mass-spectrometry. In DENV2-rep cells, we identified 228 differentially expressed host proteins (DEPs; FDR < 0.05) compared to uninfected controls (Fig. 4c). In contrast, no significant DEPs were detected in DENV2-infected cells at 24 hpi, despite comparable levels of vRNA replication and detectable changes in host transcripts (Fig. 2a, 3a). This lack of protein-level response is consistent with a global translational shutdown during acute infection and aligns with previous studies reporting minimal proteomic changes even at later timepoints of DENV2 infection in Huh7 cells[36]. These findings suggest that in acutely infected cells, transcriptional responses are not efficiently translated into changes in protein abundance, likely due to reduced translation efficiency. By contrast, DENV2-rep cells preserve active translation, allowing for dynamic adjustment of protein levels in response to persistent noncytopathic vRNA replication.

### DENV2-rep cells coordinate transcriptional and proteomic remodeling across metabolic and secretory pathways

To test the hypothesis that preserved translation in DENV2-rep cells enables coordinated transcriptional and proteomic remodeling during persistent RNA replication, we next compared the changes in mRNA and protein levels per gene. As expected, mRNA and protein fold-changes were better correlated in DENV2-rep than infected cells (Pearson’s r = 0.60 and 0.13, respectively) (Fig. 5a). These findings suggest that proteome remodeling in replicon cells is not only permitted but actively coordinated, whereas reduced translation uncouples transcriptional changes from proteomic output in acute infection.

**Figure 5.**
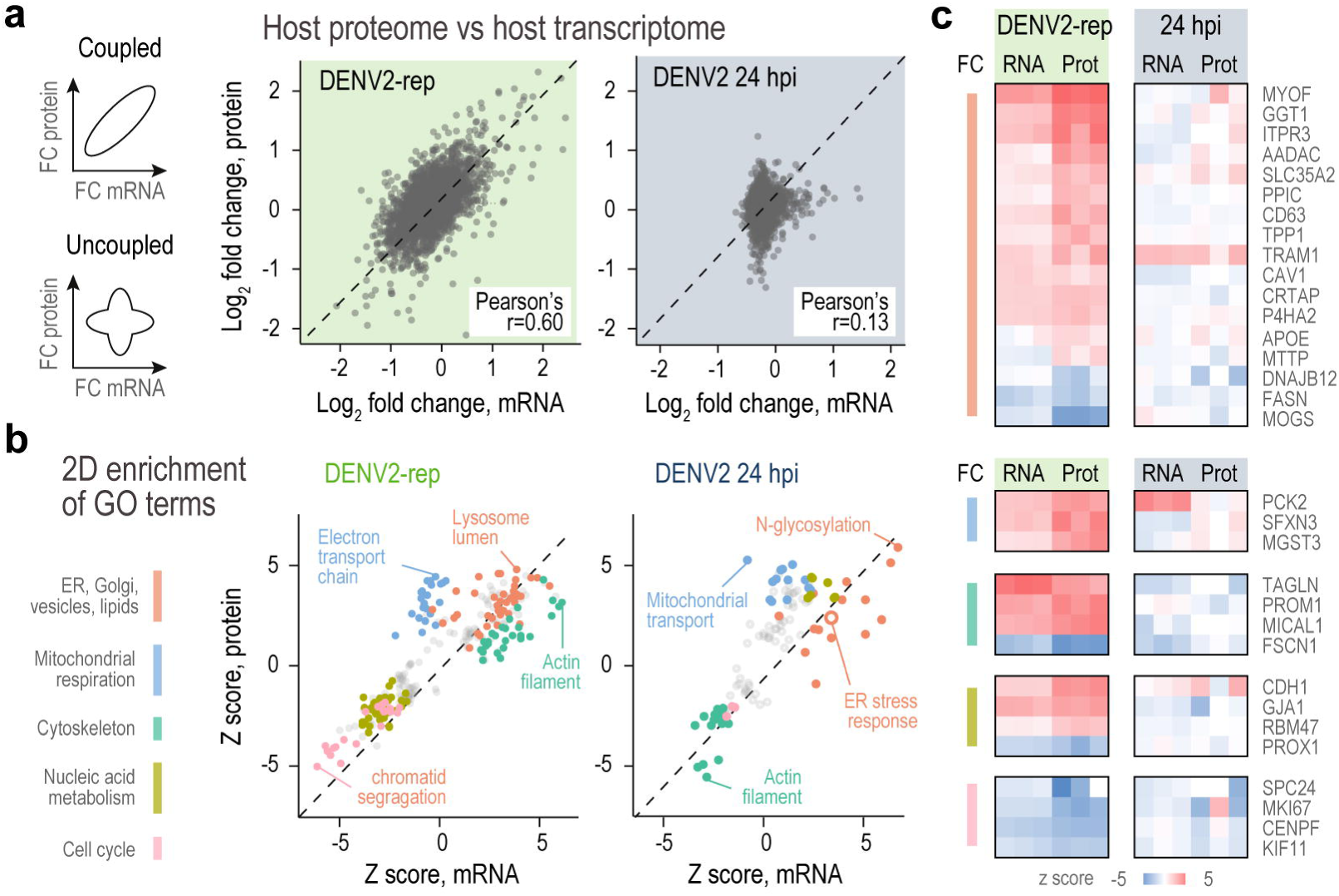
Replicon cells coordinate transcriptional and proteomic remodeling across biosynthetic pathways. (a) Correlation of mRNA and protein fold-changes per gene. Replicon cells show strong correlation (r = 0.60), while infection shows poor correlation (r = 0.13). FC, fold change. (b) Two-dimensional (2D) enrichment analysis of gene ontology (GO) categories. Replicon cells upregulate pathways spanning ER, Golgi, lysosomes, vesicular trafficking, lipid metabolism, and cytoskeleton; infection enriches for ER stress response and downregulation of cytoskeleton genes. (c) Heatmap of representative genes and proteins strongly induced or depleted in replicon cells, including lipid metabolism (GGT1, AADAC, MYOF) and ER-associated protein folding and quality-control factors (TRAM1, PDIA4, PPIC, DNAJB12, MOGS).

To identify biological pathways remodeled under each condition, we performed two-dimensional functional enrichment analysis, a method developed to identify Gene Ontology (GO) terms that vary at the mRNA and protein levels[37]. Compared to functional enrichment analysis, this approach does not involve any statistical or fold change cutoff and is less sensitive to the magnitude of gene expression changes. To account for bulk dilution effects discussed above, we used z-scored changes. In DENV2-rep cells, we observed concordant upregulation of genes involved in membrane-bound organelles and vesicular trafficking, specifically across the ER, Golgi, lysosomes, and endosomes (Fig. 5b). These organelles are intimately involved in viral polyprotein processing and replication complex formation, and are often remodeled during flavivirus infection[5,38], including persistent Dengue infection of mosquito cells[25]. Indeed, pathways involving the same organelles were also induced in acutely infected cells, although the term “ER stress response” was only elevated in infected cells (Fig. 5b).

Other categories showed concordant changes that diverged between DENV2-rep and infected cells. For example, while infected cells showed a decrease in cytoskeletal genes at both the mRNA and protein levels, DENV2-rep showed a concordant increase (Fig. 5b). Indeed, Dengue infection involves the cytoskeleton[5,38]. Interestingly, mitochondrial respiration genes were upregulated mostly at the protein level in both acute and persistent replication, suggesting post-transcriptional regulation of energy metabolism. This is in agreement with a recent report that DENV suppresses mitophagy[39].

In DENV2-rep cells, the most strongly remodeled genes at both the mRNA and protein levels included lipid metabolism and trafficking factors (GGT1, AADAC, MYOF) and ER-associated proteins involved in translocation, folding, and quality control (TRAM1, PDIA4, PPIC, DNAJB12, MOGS) (Fig. 5c). Notably, MOGS (Glucosidase I), required for NS1 maturation and whose inhibition protects against lethal DENV infection [40], was strongly suppressed in replicon cells, underscoring the therapeutic relevance of pathways remodeled during noncytopathic replication. These coordinated changes were largely absent or attenuated in acutely infected cells, highlighting the distinct host rewiring that enables persistent RNA replication while avoiding cytotoxic stress.

### Selective protein turnover limits accumulation of the viral polymerase in DENV2-rep cells

We next asked whether viral protein abundance may be subject to post-translational regulation. Despite overall similarity in viral polyprotein abundance between infection and replicon states (Fig. 6a), peptide-level mapping revealed a clear difference: NS3, the viral protease, accumulated to comparable levels under both conditions, whereas NS5 polymerase was markedly reduced in DENV2-rep cells (Fig. 6b-c). NS1, a secreted glycoprotein, also accumulated similarly across conditions, while coverage for other nonstructural proteins was insufficient for quantitative comparison (>3 peptides; Table S2).

**Figure 6.**
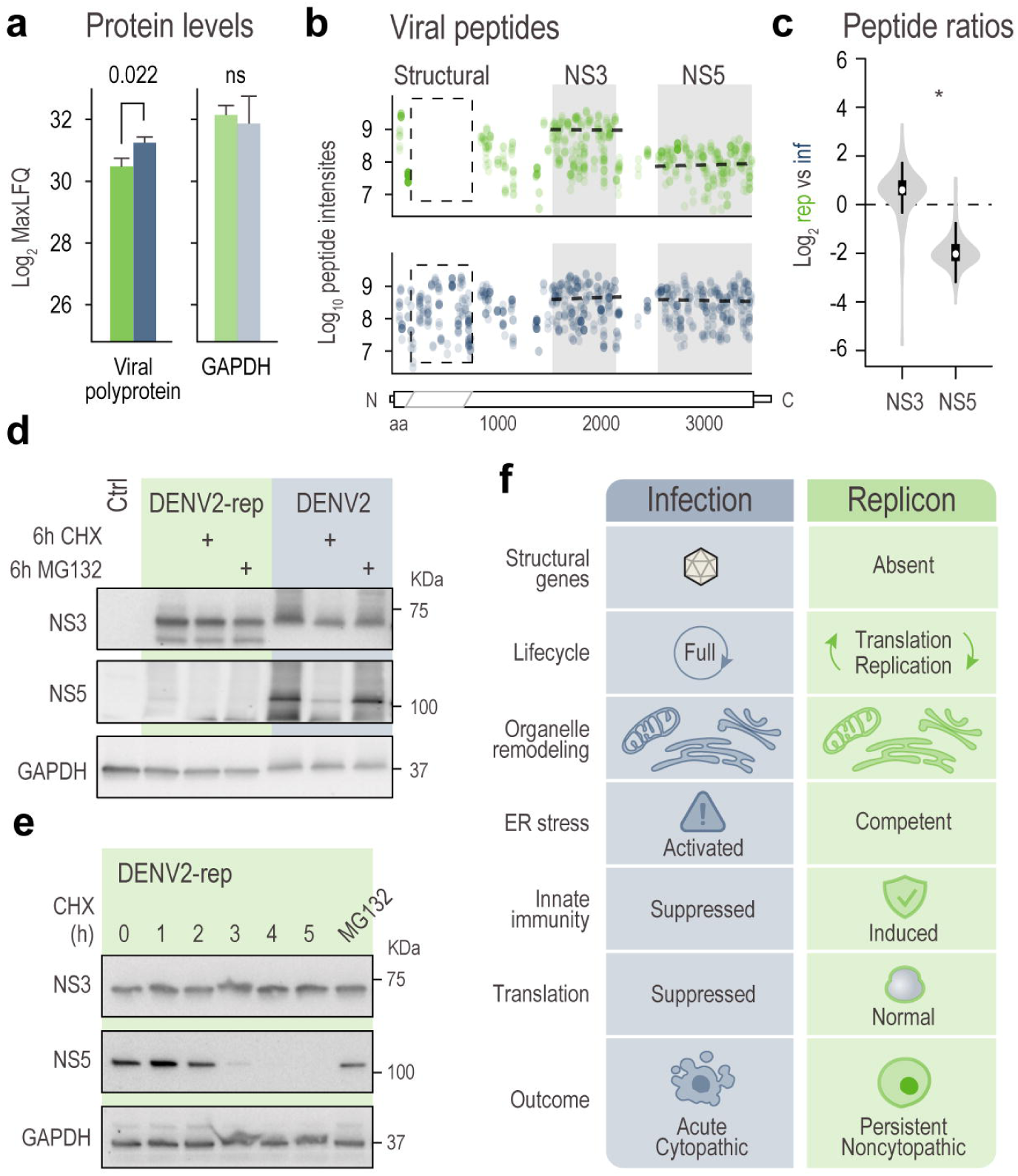
Selective protein turnover limits accumulation of the viral polymerase NS5 in DENV2-rep cells. (a) Mass spectrometry of total intracellular protein detects comparable levels of viral polyprotein in replicon and infected cells. (b-c) Peptide-level quantification reveals NS3 accumulation in both conditions, while NS5 peptides are markedly reduced in replicon cells. *, Mann Whitney p-value = 1.36x10^-75^. (d) Whole cell immunoblots confirm comparable NS3 but lower NS5 in replicon cells. Cycloheximide (CHX) reduces NS5 in infection, and proteasome inhibitor MG132 does not restore NS5 in either condition. Blots representative of 3 independent repeats. (e) Time course immunoblots following CHX treatment. NS3 and GAPDH remain stable, whereas NS5 rapidly decays in replicon cells, falling below detection within 4 hours. Blots representative of 3 independent repeats. (f) Model summarizing the divergent host responses and viral protein dynamics in infection versus replicon. Acute infection remodels organelles, induces ER stress, translation shutdown, and transcriptome/proteome uncoupling, while replicon cells remodel organelles but preserve translation and mount an antiviral response, resulting in persistent noncytopathic infection.

Immunoblotting confirmed this pattern: NS3 protein levels were comparable, but NS5 was below detection in replicon cells (Fig. 6d). Because all DENV proteins are translated as a single polyprotein, these results point to selective post-translational depletion of NS5. To test whether protein synthesis or degradation contributes to this imbalance, we treated cells with cycloheximide (CHX) or the proteasome inhibitor MG132. CHX reduced NS5 levels during acute infection, but MG132 did not increase them in either condition (Fig. 6d), suggesting that NS5 turnover is not rescued by proteasome inhibition.

We next focused our analysis on NS5 levels in DENV2-rep, which was detectable with higher loading. Following CHX treatment to block protein synthesis, NS3 and GAPDH remained stable for at least six hours, whereas NS5 declined rapidly and fell below detection within four hours (Fig. 6e). NS5 levels did not increase under MG132. Given that NS5 was also cleared more rapidly than NS3 in acutely infected cells (Fig. 6d), this could represent a previously unrecognized vulnerability of dengue replication, highlighting NS5 stability as a potential therapeutic target.

Together, these findings establish that the replicon state is characterized by depletion of the viral polymerase NS5, even in the presence of robust RNA replication and polyprotein synthesis. This reveals an additional layer of viral regulation in which protein turnover, rather than translation alone, sets the abundance of key replication enzymes. This uncovers a new layer of regulation in the viral life cycle, one in which protein stability, not just translation, limits the abundance of key viral enzymes. It also highlights a potential vulnerability of noncytopathic RNA replication that could be exploited therapeutically.

## Discussion

RNA viruses profoundly rewire the intracellular environment to favor their replication, often at the expense of host cell viability [5]. While cytopathic infection models have historically shaped our understanding of these processes, noncytopathic systems such as viral replicons offer a complementary view, isolating the effects of RNA translation and replication from entry, assembly, and egress. Here, we show that a stable DENV2 replicon (DENV2-rep) establishes a distinct host cell state that accommodates long-term viral RNA replication without triggering the cytopathic stress responses observed during acute infection. Our model (Fig. 6f) highlights how acute infection and persistent replicon replication diverge at multiple checkpoints, from proteostasis to innate immune signaling, providing an initial framework for understanding persistence.

Our findings help resolve uncertainties about ER stress in flavivirus infection. Prior studies showed that live DENV2 infection triggers ER stress, activates the UPR, and suppresses global translation—responses conserved across Flaviviridae[41] and even in an HCV replicon model [42]. In contrast, DENV2-rep cells supported long-term replication with preserved translation, intact polysomes, and minimal UPR induction, indicating that structural protein expression—rather than RNA replication alone—is the primary driver of ER stress and cytotoxicity (Fig. 6f).

CRISPR screens using DENV2-rep cells identified proviral factors involved in ER protein folding [20], and our data show that some ER-related genes are selectively regulated, suggesting that replicon cells establish a distinct steady-state equilibrium that permits continued proliferation despite persistent RNA replication. This equilibrium differs from both acutely infected and uninfected cells, reflecting active remodeling of biosynthetic and metabolic pathways. By comparing infection and replicon states, we clarify that nonstructural proteins are sufficient to remodel the ER, Golgi, lysosomal, and lipid metabolic networks—features emphasized in multiple flavivirus studies[41]—but without inducing PERK/ATF4 signaling or global translational arrest. Instead, biosynthetic activity remains intact, enabling selective remodeling of the cellular landscape.

Unexpectedly, DENV2-rep cells mounted an antiviral transcriptional program, including induction of interferon-stimulated genes (ISGs), and resisted superinfection. This is striking given the RIG-I deficiency of Huh7 cells [43]. Whereas acute infection suppressed ISG induction, replicon cells upregulated ISGs such as OASL, IFIT2, MX1, and RSAD2, consistent with reports from dengue patients and vaccine recipients [30,31]. Another key distinction was the preservation of translation and the coupling of transcriptomic and proteomic remodeling in replicon cells. While acute infection uncoupled mRNA and protein changes, replicon cells maintained tight mRNA-protein correlations, enabling remodeling of metabolic, cytoskeletal, and vesicular trafficking pathways. Similar organellar remodeling was observed with persistent Dengue infection of mosquito cells[25]. Such coordination provides a mechanistic explanation for how cells can adapt to persistent RNA replication while avoiding cytotoxicity.

Finally, our study uncovered marked post-translational turnover of NS5, the viral polymerase, in replicon cells. Despite equimolar expression of all nonstructural proteins from the viral polyprotein, NS5 was depleted relative to NS3 and NS1. This was not explained by RNA levels, translation efficiency, or proteasome activity. NS5 turnover was also detectable during acute infection, but less pronounced, suggesting stabilization by higher RNA load or structural protein interactions. NS5 is known to shuttle between cytoplasm and nucleus in other flaviviruses, where nuclear trafficking influences immune modulation and host transcription. [27,44–46]Its selective depletion in replicon cells suggests an additional layer of regulation in which viral polymerase stability, not just translation, constrains persistent replication. Such turnover may represent a host defense strategy or, alternatively, a therapeutic vulnerability.

In summary, the DENV2 replicon clarifies which features of Flaviviridae–host interactions arise from RNA replication alone versus structural protein expression. While replicons do not recapitulate all aspects of infection, they uniquely reveal how cells can tolerate long-term viral RNA replication with preserved translation, selective organelle remodeling, induction of antiviral defenses, and post-translational regulation of viral proteins. This framework complements existing cytopathic models, highlights candidate pathways for host-directed antiviral strategies, and positions replicons as powerful tools for dissecting long-term virus–host interactions in a controlled, noncytopathic context.

## Methods

### Cell lines, culture conditions and viruses

Human hepatocyte-derived carcinoma Huh7.5.1 cells were sourced from Apath (Dr. Charles M. Rice) and Scripps Research (Dr. Francis V. Chisari), and were maintained in Dulbecco’s Modified Eagle Medium (DMEM) F-12 mixture (DMEM/F-12) supplemented with 10% fetal bovine serum (FBS), 1× non-essential amino acids, and penicillin/streptomycin at 37°C in a humidified 5% CO incubator. DENV2 replicon cell lines (DENV2-rep) were generated and maintained under Zeocin (InvivoGen) or Blasticidin (Thermo Fisher) selection as indicated. The infectious dengue virus type 2 (DENV2) clone 16681 and dengue type 2 expressing *Renilla* luciferase (DENV2-Luc) were a gift from Jan Carette (Stanford University).

### Generation of stable DENV2 Replicon (DENV2-rep) cell line

DENV2 replicon constructs were derived from the DENV2 16681 infectious clone. To create the DENV2-rep plasmid, we replaced the first 2,876 nucleotides of the viral genome with a gene synthesis product (Integrated DNA Technologies) containing *Sac*I, T7 promoter, full-length DENV2 5’UTR and first 102 nucleotides of the coding region, EGFP, a PEST degradation domain, Blasticidin and an F2A ribosome skip site. This is followed in-frame by the last 72 nucleotides of Envelope (E) and the start of NS1 up to an endogenous *Eco*RI site. Blasticidin was replaced with Zeocin resistance cassette by overlap PCR and ligation. For *in vitro* transcription of DENV-2 replicon RNA, the DENV-2 replicon plasmid was linearized with *Xba*I. 1 µg of linearized DENV-2 replicon DNA was transcribed into RNA using a MEGAscript T7 Transcription Kit (Ambion, AM1334) with the addition of 4 mM of m^7^G(5’)ppp(5’)G cap analog (NEB, S1405S) per reaction. The reaction was incubated for 4 h at 33°C and then treated with Turbo DNAse at 37°C for 15 min. Replicon RNA was purified using an RNA Clean and Concentrator kit (Zymo Research, R1017) and stored at -80°C. Huh7.5.1-cas9 cells were seeded in a 12-well plate (200,000 cells per well) for transfection with purified replicon RNA the following day. Next, 1 µg of purified replicon RNA was combined with an mRNA transfection reagent (Mirus, MIR2225) in accordance with the manufacturer’s protocol. The transfected cells were visualized under a fluorescence EVOS microscope 24 h after transfection for the appearance of eGFP-expressing cells. Cells were placed under antibiotic selection (4 µg/mL blasticidin; 200 µg/mL Zeocin) 2 d after RNA transfection. Cells were expanded and frozen.

### Virus infections

Huh7.5.1 or DENV2-rep cells were plated in either 96-or 12-well plates at a density of 0.8x10^4^ or 1x10^5^ cells/well, respectively. The following day, cultures were infected with wild-type DENV2 16681 or DENV2-Luc at MOI = 0.1 or 5, as follows. Cells were washed once with PBS and viral inoculum was added into fresh DMEM/F-12 media supplemented with 2% FBS. Cells were incubated with the inoculum at 37°C for 2 hours with occasional shaking. After 2 hours, inoculum was replaced with fresh DMEM/F-12 media supplemented with 10% FBS.

### Immunofluorescent microscopy

Huh7.5.1-Cas9 (parental) and Huh7.5.1-DENV-replicon (eGFP-blast) cells were seeded at a density of 10,000 cells per well in a glass-bottom 96-well plates coated with 50 µg/mL collagen (Gibco, A1048401). The following day, Huh7.5.1-Cas9 cells were infected with DENV2 at an MOI of 2. At 24 hours post-infection, cells were fixed in 4% paraformaldehyde in PBS for 15 minutes at room temperature and permeabilized with 0.2% Triton X-100 in PBS for 10 minutes. Permeabilized cells were then blocked for 1 hour at room temperature in blocking buffer (3% BSA, 0.1% Triton X-100, and 0.1% sodium azide in PBS). Primary antibody (Anti dsRNA monoclonal J2, Jena Biosciences RNT-SCI-10010200) was diluted in blocking buffer (1:1000 dilution) and incubated overnight at 4C. Secondary antibody (Goat anti-mouse AF-647, Thermo Fisher Scientific A-21235) was diluted in blocking buffer (1:1000 dilution) and incubated for 1 hour at room temperature. Cells were washed 3X in PBS then imaged using a Leica DMi8 inverted confocal microscope equipped with a 63× oil-immersion objective (NA 1.47) and controlled by Micro-Manager 2.0 software.

### Flow cytometry

DENV2-rep and parental control cells were assayed for EGFP expression by flow cytometry (Beckman Coulter CytoFlex) and analyzed using FlowJo software v 9. For cell cycle analysis, DNA was stained by adding Hoechst 33342 (Thermo Fisher) directly into culture media at a final concentration of 5 ng/mL for 1 hour at 37°C. A minimum of 10,000 cells were analyzed per sample.

### Transcriptomics (RNA-Seq)

Huh7.5.1-Cas9 (parental) and Huh7.5.1-DENV-replicon cells were seeded in triplicate at a density of 200,000 cells per well in 6-well plates. The following day, Huh7.5.1-Cas9 cells were infected with DENV2 at an MOI = 5. At 24 hpi, total RNA was extracted from uninfected, infected, and DENV-replicon cells using the Zymo Quick-RNA Miniprep Plus Kit (Zymo Research, R1057), and samples were stored at -80°C until further processing. For RNA sequencing, 100 ng of purified RNA from each sample was used as input for library preparation. An ERCC spike-in mix (External RNA Controls Consortium) was added to each sample as an internal positive control. To deplete ribosomal RNA from mammalian transcripts, FastSelect-rRNA HMR (Qiagen) was included at a 1:10 dilution. Libraries were prepared using the NEBNext Ultra II Directional RNA Library Prep Kit (New England Biolabs, E7760S), following the manufacturer’s protocol. Briefly, RNA was reverse-transcribed into cDNA, which was then used for construction and barcoding of sequencing libraries. Final libraries were sequenced using 146-nucleotide paired-end reads on the Illumina NextSeq 2000 platform (P3 flow cell), with a target sequencing depth of at least 50 million reads per sample. After sequencing was completed, reads from each sample were mapped to the human GRCh38 genome concatenated to the ERCC sequences and the DENV2-16681 genome using STAR Aligner[47] and counted using htseq-count[48]. Sequencing depth was estimated using SAMtools-depth[49]. Differential gene expression analysis was performed using custom R scripts based on DESeq2[50]. GSEA was performed using WebGestalt[51].

### qPCR

Cell lysates harvested from DENV-infected and control samples were analyzed by RT-qPCR using the Cells-to-Ct kit (Thermo Fisher Scientific, A25600) in accordance with manufacturer’s protocol for qPCR readout of vRNA relative to 18S RNA (housekeeping gene). The following qPCR primers were used: universal DENV forward, 5’-GGTTAGAGGAGACCCCTCCC-3’; universal DENV reverse, 5’-GGTCTCCTCTAACCTCTAGTCC-3’; 18S forward, 5’-AGAAACGGCTACCACATCCA-3’; 18S reverse: 5’-CACCAGACTTGCCCTCCA-3’; GAPDH forward, 5’-AGGTCGGAGTCAACGGAT-3’; and GAPDH reverse, 5’-TCCTGGAAGATGGTGATG-3’.

Raw Ct values were extracted and normalized to 18S or GAPDH for further analysis.

### Luciferase

Cells infected with DENV-2-Luc were washed three times with PBS and lysed. Then *Renilla* luciferase assay reagent (Promega) was added to each well in accordance with the manufacturer’s protocol. Luciferase signal was measured using luminescence readout per well on the Spectramax i3x plate reader.

### SDS-PAGE and immunoblotting

For immunoblotting, adherent cells were washed twice with ice-cold PBS and lysed on plate with RIPA buffer (25 mM Tris-HCl pH=7.5, 150 mM NaCl, 1% NP-40, 0.5% Sodium deoxycholate) supplemented with 1 mM DTT, Complete EDTA-free protease inhibitor cocktail, and 50 units/mL benzonase (Millipore Sigma) to remove DNA. Lysis was performed on ice for 20 min and lysates were clarified by centrifugation for 10 min at 12,000 x g, 4°C. To label nascent chains for puromycin immunoblotting, 1 µM puromycin (Thermo Fisher) was added to tissue culture media for 10 min at 37°C before harvesting. Protein concentration was determined by BCA assay (Thermo Fisher) and reconstituted in 1x Laemmli sample buffer (Bio-Rad) supplemented with fresh 1% 2-mercaptoethanol. 10-20 µg of each sample was resolved on 4-20% denaturing gels (Bio-Rad) and transferred to 0.2 µm PVDF membranes using a wet transfer apparatus. Membranes were blocked with 4% molecular biology grade BSA (Millipore Sigma) in tris-buffered saline supplemented with 0.1% Tween-20 (Millipore Sigma, TBST) for 1 h at RT then probed with primary antibodies for 2 h at RT. Primary antibodies (puromycin, Millipore MABE343; ATF4 Cell Signaling 11815; DDIT3/CHOP Cell Signaling 5554; GAPDH Genetex GTX627408; DENV2 NS3 Genetex GTX629477; DENV2 NS5 Genetex GTX629447) were diluted 1:1000 in 4% BSA/TBST supplemented with 0.02% sodium azide. Secondary antibodies were diluted 1:10,000 in TBST. Western blot detection was done using ECL Plus Western Blotting Substrate (Thermo Fisher) and images were taken either by film radiography or BioRad GelDoc imager. Densitometry analysis was performed using ImageJ version 1.52a.

### Polysome profiles using sucrose gradients

Cells were harvested by scraping in ice-cold PBS (with calcium and magnesium, Thermo Fisher), centrifuged 1000 x g for 5 min at 4°C, resuspended in PBS and centrifuged again. Cell pellets were flash frozen in liquid nitrogen. On the day of the experiment, pellets were thawed on ice and resuspended in 200 µl polysome buffer (25 mM Tris-HCl pH=7.5, 150 mM NaCl, 10 mM MgCl_2_, 2 mM dithiothreitol (DTT) and Complete EDTA-free protease inhibitor cocktail (Roche)). Triton X-100 and sodium deoxycholate (Millipore Sigma) were added to a final concentration of 1%. After 20 min on ice, the samples were centrifuged at 20,000 x g for 10 min at 4°C to remove cell debris. Clarified lysates were loaded on 10-50% sucrose gradients in polysome buffer and subjected to ultracentrifugation at 41,000 rpm in an SW41.Ti swinging bucket rotor (Beckman Coulter) for 150 min at 4°C. Equal volume fractions were collected using Gradient Station (BioComp) with continuous monitoring of rRNA at UV254.

### LC-MS/MS sample preparation and data acquisition

Cells were lysed in urea buffer and protein concentration was determined by BCA assay (Thermo Fisher). 1-2 µg total protein were subjected to reduction and alkylation by incubation with 10 mM DTT (Thermo Fisher) for 1 h at room temperature followed by 5 mM iodoacetamide (Thermo Fisher) for 1h at room temperature, in the dark. Samples were adjusted to 2M urea in 25 mM ammonium bicarbonate (pH 7.5) and sequencing-grade modified trypsin and lys-C (Promega) were added at 1:50 enzyme to protein ratio. Digests were incubated overnight at 25°C and acidified with 0.1% trifluoroacetic acid. Peptides were desalted with in-house styrenedivinylbenzene reversed phase sulfonate packed stagetips, dried and resuspended in 2% acetonitrile/0.1% TFA. Peptides were analyzed on a Fusion Lumos mass spectrometer (Thermo Fisher) equipped with a Thermo EASY-nLC 1200 LC system (Thermo Fisher). Peptides were separated by capillary reverse phase chromatography on a 25 cm column (75 µm inner diameter, packed with 1.7 µm C18 resin, AUR3-25075C18, Ionopticks, Victoria Australia). Peptides were introduced using a two-step linear gradient with 3-27 % buffer B (0.1% (v/v) formic acid in 80% (v/v) acetonitrile) for 52.5 min followed by 27-40 % buffer B for 14.5 min at a flow rate of 300 nL/min. Column temperature was maintained at 50°C throughout the procedure. Data was acquired in top speed data dependent mode with a duty cycle time of 1 s. Full MS scans were acquired in the Orbitrap mass analyzer (FTMS) with a resolution of 120,000 (FWHM) and m/z scan range of 375-1500 m/z. Selected precursor ions were subjected to fragmentation using higher-energy collisional dissociation (HCD) with quadrupole isolation window of 0.7 m/z, and normalized collision energy of 31%. HCD fragments were analyzed in the Ion Trap mass analyzer (ITMS) set to Turbo scan rate. Fragmented ions were dynamically excluded from further selection for a period of 60 seconds. The AGC target was set to 1,000,000 and 10,000 for full FTMS and ITMS scans, respectively. The maximum injection time was set to Auto for both full FTMS scans and ITMS scans.

### MS data processing

Raw MS data for polysomes were processed using MSFragger v3.8, FragPipe v20.0, IonQuant v1.9.8 and Philosopher v5.0.0. MS/MS spectra were searched against the forward and reverse human proteome (UP000005640) and type 2 DENV (P29990). Cysteine carbamidomethylation was chosen as fixed modification and methionine oxidation and STY phosphorylation as variable modifications. Precursor and fragment mass tolerances were set at 4.5 and 20 ppm, respectively. Maximum allowed false discovery rate (FDR) was <0.01 at both the peptide and protein levels, based on a standard target-decoy database approach. Match between runs was enabled. Default settings were used for LFQ workflow. MaxLFQ intensities were extracted from Combined_protein.tsv and used for downstream analysis using Perseus 2.0.7.0. Non-normalized intensities were extracted from Combined_peptide.tsv to analyze the distribution of dengue peptides and the per-peptide ratios between infected and DENV2-rep cells. ENSG IDs were used for integration with RNA-seq data.

## Acknowledgements

We thank the CZB-SF Genomics platform lead Norma Neff and team members Angela Detweiler, Honey Mekonen, Sheryl Paul, and Amanda Seng for consultation and technical support for library preparation and sequencing throughout this study. This study was funded by the Chan Zuckerberg Initiative.

## Author contributions

K.C., D.K., K.F.W. designed and performed experiments and analyzed data. F.M. acquired LC-MS/MS data. J.E.E. supervised and provided funding for mass-spectrometry. R.A. and A.K. conceived the study. R.A. designed and performed experiments, analyzed data and wrote the manuscript.

